# RUNIMC: An R-based package for imaging mass cytometry data analysis and pipeline validation

**DOI:** 10.1101/2021.09.14.460258

**Authors:** Luigi Dolcetti, Paul R Barber, Gregory Weitsman, Selvam Thavaraj, Kenrick Ng, Julie Nuo En Chan, Piers Patten, Rami Mustapha, Jinhai Deng, Tony Ng

**Affiliations:** Richard Dimbleby Laboratory of Cancer Research, School of Cancer & Pharmaceutical Sciences, King’s College London, London, UK; UCL Cancer Institute, Paul O’Gorman Building, University College London, London, UK; Comprehensive Cancer Centre, School of Cancer & Pharmaceutical Sciences, King’s College London, London, UK; Head & Neck Pathology, Guy’s and St Thomas’ NHS Foundation Trust, London, UK; Faculty of Oral, Dental and Craniofacial Science, King’s College London, London, UK; Department of Haematology, King’s College Hospital, London UK; Comprehensive Cancer Centre, Faculty of Life Sciences and Medicine, King’s College London, London, UK; Breast Cancer Now Research Unit, School of Cancer & Pharmaceutical Sciences, King’s College London, London, UK

## Abstract

We propose a novel pipeline for the analysis of imaging mass cytometry data, comparing an unbiased approach, representing the actual gold standard, with a novel biased method. We made use of both synthetic/ controlled datasets as well as two datasets obtained from FFPE sections of follicular lymphoma, and head and neck patients, stained with a 14 and 29-markers panels respectively. The novel pipeline, denominated RUNIMC, has been completely developed in R and contained in a single package. The novelty resides in the ease with which multi-class random forest classifier can be used to classify image features, making the pathologist’s and expert classification pivotal, and the use of a random forest regression approach that permits a better detection of cell boundaries, and alleviates the necessity of relying on a perfect nuclear staining.

## INTRODUCTION

Imaging in cancer is probably one amongst the major contributors to the reduction of tumour related mortality^1^ and, by way of the so called molecular imaging^2^ and hybrid staining^3^, a source of high-content data for basic science. Microscopy in particular, as well as many other techniques in life science, has been subjected, in recent years, to a progressive dimensionality inflation, as a consequence of technological improvements such as the use of metal tagged antibodies and correlated detection systems, i.e. imaging mass cytometry (IMC) and multiplexed ion beam imaging (MIBI)^4^, that required the development of new data analysis tools and visualisation approaches^5^. Nonetheless, when the subject of imaging is an animal or patient tissue, an essential step is the identification of subunits, most commonly single cells, that can help to disentangle data complexity. Cell biologist and pathologist can rely on a large choice of tools spanning the full spectrum of free, open-source, and proprietary software on one hand, and generalist or high specialized applications on the other. Image-J^6^ (and its direct derivative Fiji^7^) is one of the most established generalist container that offer invaluable flexibility due to its architecture based on plugins and macros implementations. Image-j offers numerous cell segmentation options mainly based on watershed and thresholding algorithms. CellProfiler^8^, in the context of IMC, gained great popularity for the easy multistep segmentation approach it takes and the possibility to manually fine-tune segmentation parameters. Thresholding and clustering methods suffer from several limitations when pixel intensity histograms are not well equalized or in a situation of elevated noise. Slope difference distribution such as in Wang’s method^9^ overcome some of these issues although still lacking a full unsupervised implementation^10^. A considerable improvement over cell segmentation based on direct thresholding of the acquired image, has been the introduction of machine-learning approaches, such as that implemented in ilastik^11^, that permits to feed to the segmentation algorithm one metric that is an estimation, based on the information conveyed by multiple images or layers, of the probability of a pixel to belong to a certain category or cluster, or the use of deep-learning algoritms^12^, topic that will be not expanded in this introduction because it is beyond the purpose of the present article.

The approach we are presenting in this paper (and we used recently in practice in Barber et al., submitted paper), implemented as a new self-contained R-based package denominated “RUNIMC”, moves from the use of machine-learning classification and extends it with regard to two main aspects. First, the classification process is conducted on multiple classes, for example non-overlapping cell lineages (T-lymphocytes, B-lymphocytes, macrophages, etc), and use these classes to create a pre-segmentation partition of the image. Second, within each partition, a prediction of the position of each pixel within a putative cell is calculated on the basis of a second random forest regression model. The predicted position can then be used to model cell segmentation by means of different approaches. In particular for this paper, we implemented an algorithm that evaluates areas as the local expansion of seed-pixels, but different algorithm (also included in the package) have been tried such as slope evaluation along radii extending from a centroid pixel, and geometrical constrained segmentation.

Validation is a problem with no gold standard, and usually solved by “does the data look sensible?” approach. We propose the use of simulated data based on common general marker distribution patterns (membranous, cytoplasmic etc.) and common cell shapes (round, protrusions, etc.) with variations simulated by randomisation of shape forming parameters and the addition of Gaussian pixel noise. This companion package, called “barbieHistologist” to reflect to the modelling nature of this element, works with RUNIMC to generate data with a known gold standard segmentation with which to compare segmentation pipelines. We have used this to objectively compare the segmentations of a “reference” pipeline with our novel “alternative” pipeline.

## MATERIAL AND METHODS

### FFPE histology staining

Five µm FFPE histological slides from head and neck cancer patients and follicular lymphoma were stained with a panel of metal conjugated antibody as detailed in figure 1C. Where conjugated antibodies for specific targets were not available, metal conjugation was performed with Maxpar X8 antibody labelling kit (Fluidigm) following manufacturer instructions.

**Figure 1:**
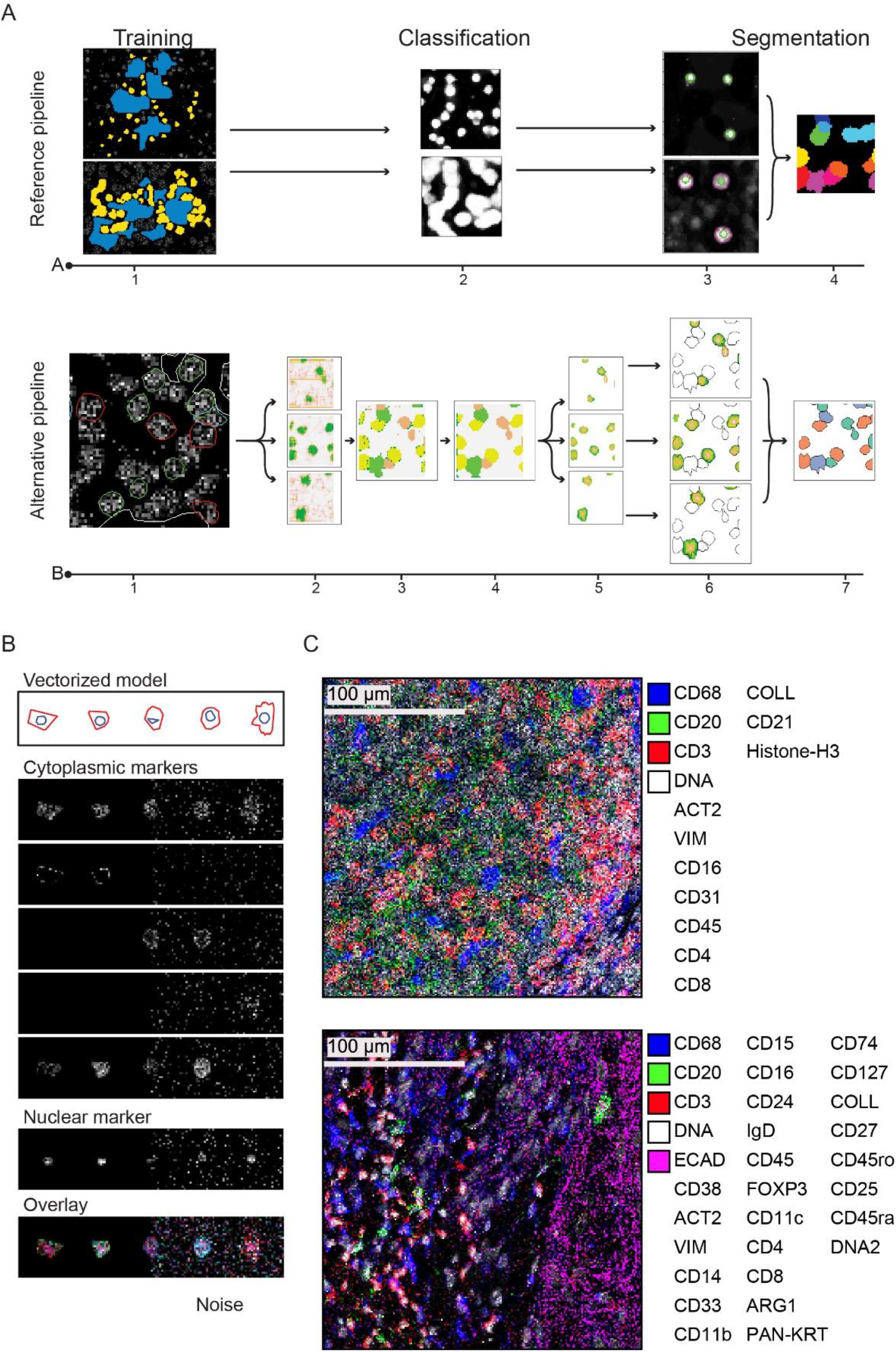
pipeline comparison and methods. Schematic outline of the pipeline comparison (A). The *reference* pipeline (upper row) and the novel *alternative* pipeline (lower row) both start with a manual annotation of the image that permits the extraction of training features (1). Annotation, training, and class prediction (2) for *reference* has been carried out using ilastik software in two steps, using separately the nuclear staining to identify nuclei and all the other channels to identify cell cytoplasm. Annotation for *alternative* has been carried out using a Shiny-app GUI front end included in the RUNIMC package identifying multiple classes on the basis of lineage markers (for example CD4^+^, CD8^+^, CD20^+^, etc. cells). In the case of *alternative*, classification is a multi-step process including cell level classification (2-3), an optional local clean-up (4), sub-cell regression (5). *Reference* segmentation (3) was accomplished using CellProfiler in two steps, segmenting nuclei first as primary object (upper row) and identifying single cells (secondary object) as nucleus area expansion and deconvolving cell clusters via the implemented Otsu algorithm (3-4). *Alternative* segmentation is carried out analyzing the gradient generated in the regression step (6) and polygons are aggregated in a final step (7). Example of synthetic image generation (B). In order to compare the two pipelines against a controlled test, synthetic data has been generated with the R package barbieHistologist. Cell outlines are parametrically generated (a) and associated to simulated markers that can be either membranal, cytoplasmatic or nuclear. The lower picture represents color coded overlay of the synthetic markers above and the addition of random noise (right portion each image). Representative image from two case studies used for the comparison(C). FFPE histology of lymph nodes from follicular lymphoma patient stained with a panel of antibody listed on the right (upper panel), and FFPE histology of head and neck tumor patient (lower panel).

Antigen retrieval was performed on a Ventana Bench Mark Ultra with CC1 buffer (Roche, 950-224). Slides were blocked for 1 hour at RT in 5% BSA, 5 mg/ml human IgG, PBS, and stained ON at 4°C in 5% BSA, PBS. DNA counterstain was performed with Iridium (Fluidigm, 201192B) 125 nM in PBS for 30 minutes at RT.

Ablation and data acquisition were performed on a Fluidigm Hyperion at Guy’s and St Thomas’ BRC.

### Generation of synthetic data

To obtain multiple test data sets with given features, we developed an R package for the generation of synthetic data mimicking a complex of cells in a tissue or on a plate. Individual synthetic cells are defined via an S4 object containing the 2D geometrical description of different compartments (cytoplasm, nucleus, organelles) and the parametric description of specific markers that can be associated with each geometry. Prototypic shapes and markers are defined as parametric objects, i.e. geometric features such as dimension, roundness, convexity or intensity and pattern of staining are specified as mean and standard deviation, handing over to the algorithm the task of organizing a cell population with controlled phenotypical variability within a specific space and according to given proportions.

When an entire cell population is defined, each marker is impressed on a separate pixel-based raster layer with given intensity statistics (mean, sd), and filters (noise, blurring, etc.) can be applied to mimic the acquisition process from various instruments such as a Fluidigm Hyperion, Multiplexed ion beam imaging or fluorescence imaging. The outputs images from the simulation were compared to real data and assessed for noise content and cell definition (fig 1B and C).

In order to stress-test the two pipelines we set up four different scenarios that would be frequently experienced in a real analysis. First, we test the response to cell density as it would be found in different tissue areas, such as a large patch of connective tissue with a few dispersed cells or a largely and heavily congested field from a lymph node. Second, we simulated an increased rate of geometrical complexity i.e., the deviation from a round shape and the appearance of cellular protrusions, such as for dendritic cells. Third, we tested the effect of applying the two pipelines to a set of non-homogeneous samples. To achieve this comparison, we simply added a progressively heavier amount of noise to similar images, using a small portion of each image as training set. Fourth, we assessed the capacity of the two pipelines to discriminate correlated phenotypes for example lymphocytes subsets such as CD3^+^CD8^+^CCR7^-^ CD45RO^+^ effector memory from CD3^+^CD8^+^CCR7^+^CD45RO^+^ central memory, in a context of a large and repetitive tissue as it would be a tumor infiltrated by immune cells. Finally, we tested the two pipelines in real case study, producing the segmentation of 3 images taken from independent follicular lymphoma patients, and 14 ROIs derived from two head and neck cancer patients.

### Reference pipeline

As reference method for comparison with our new method, we made use of two well established tools: ilastik^11^ for pixel classification and cellProfiler^8^ for segmentation. In its most straight forward form this pipeline requires training/prediction using a random forest classifier in two separate steps for the identification of cell nuclei and, subsequently, for cell cytoplasm. The probability maps generated by means of ilastik (figure 1A step 1 and 2) have been fed to CellProfiler to identify primary objects (nuclei) that have been propagated using Otsu thresholding algorithm to identify secondary objects (complete single cells). This pipeline is an implementation of a previously published method by Schulz et al^13^.

### Alternative novel pipeline

The novel alternative pipeline we are presenting is “core to RUNIMC” and resembles, in principle, the random forest-based machine-learning adopted by the reference pipeline with few essential variations.

The pipeline, with reference to the methods contained within RUNIMC, is now described.

Fluidigm’s proprietary MCD data files are exported by means of MCD viewer (Fluidigm) to plain XYZ text files and loaded into R via the function *initStudy* that both initializes a new study, creating the basic object and file structure used later by RUNIMC, and transforms the images from a tabular format to an object of class *IMC_RasterStack* (a simple extension of a *RasterStack* object from the package *raster*^14,15^).

The images are now accessible via the function *runIMC* that opens a *Shiny* GUI frontend, the basic function of which is to enable the user to identify and sample representative cells and features that will be used to train the random forest classifier. Differently from the *reference* approach, the selection of region of interest (ROI) that define single cell areas is multiclass in principle. In other words, mutually exclusive cell-types can be aggregated into different classes: for example, CD8, CD4, CD20 lymphocytes, etc. should be annotated as belonging to different classes. This contrasts with the *reference* approach which identifies all cells in a single class. Moreover, *runIMC* takes note of geometrical parameters such as the area, perimeter and roundness of each polygon used to identify specific cells; these parameters can be used later to filter and adjust the product of segmentation or used directly in a specific segmentation algorithm (figure 1.A, step 1).

Similarly to ilastik, the dataset is enriched via the use of built in or custom non-linear transformations that are specified via the method *addFilter* and calculated by *deployFilters*, which introduce measurements of local image texture and intensity as derived features. The set of original features and derived features is then extracted from the training set via the method *extractTrainingFeatures* and fed to a random forest classifier via the method *makeClassificationModel*. This trained classifier can be used in the next steps on the complete data set to produce predictions of the locations of all cells of the trained classes

Dissimilarly from the *reference* pipeline, this novel pipeline does not pass the probability maps obtained from the random forest prediction to the segmentation method directly (figure 1.A, step 2), but instead produces an unambiguous classification map (figure 1.A, step 3) that can be subjected to a topological-aware clean-up via the method *localCorrection* (figure 1.A, step 4).

At this point a second random forest model for regression instead of classification is calculated. The position of each pixel of the training set within the polygon it belongs to has been evaluated in terms of distance from the centroid of the polygon, and the concentric layer position obtained from a process of erosion of the polygon margin. A prediction of the distance/position, deployed separately for each classification map, will form a new set of maps (figure 1.A step 5) that are going to be passed to the segmentation algorithm.

The segmentation algorithm, specified via the method *addSegmentationDirectives* and started with the command *segment*, that has been used for all the tests presented in this paper is based on the evaluation of the gradient (distance from the centroid or the position of each pixel) for growing areas in an algorithm that resemble partially a watershed and dilatation approach.

The product of segmentation as the vectorized representation of each area, together with the mean value of each marker is then stored in a *sf* data frame (package *sf*^*16*^) ready for downstream analysis or plotting.

RUNIMC contains supplementary utility function to ease the interaction with other tools, for example importing cell segmentation masks from CellProfiler, via *importTiffMask*, their vectorization using the function *lazyCatMap* and the extraction of mean or other statistics from segmented images by means of the function *extractMeanPixel*.

The complete R package is available at https://github.com/luigidolcetti/RUNIMC.git, a clean and improved version will be available soon.

The package for the creation of synthetic data is available at https://github.com/luigidolcetti/barbieHistologist.git, and the complete set of scripts that make use of the two packages is available here https://github.com/luigidolcetti/RUNIMC_paper_scripts.git.

### Metrics definition

#### Normalized Shannon entropy

We used normalized Shannon entropy to quantify the amount of incorrect information generated by the process of segmentation. In brief, the vectorized model generated with the package barbieHistologist, is transformed in a bitmap image (raster matrix), where each pixel will receive the index of the cell by which is covered, this constitutes the ground truth (GT). The same model is used to generate the stack of images (one image per marker) that will be analyzed.

After a stack of images has been analyzed with both pipelines, the matrix representing the GT is used to identify the content, in terms of pixels, of each segmented area. One area would, ideally, contain a homogeneous population of pixel marked with the same GT index, otherwise a mixture of labels with different proportions would contribute to increase the associated entropy, H.

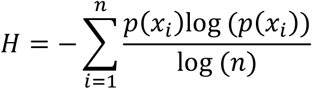

Where *p*(*x*_*i*_) is the probability of the label *i* among *n* labels contained in each area.

If the area is perfectly homogeneous, H will be zero, with larger H resulting from a mixture of labels.

#### Intersection over union

In order to quantify how well the empirical segmentation matched the GT, we evaluated for each cell the intersection over union (IOU) of the empirical polygon and the GT polygon, expressed as:

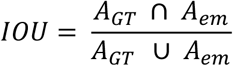

Where *A*_*GT*_ is the area of the GT polygon and *A*_*em*_ is the area generated by the segmentation.

The most common pixel label inside each polygon dictated the correspondence between predicted cell and GT cell. If the segmentation matches the GT then the IOU will be 1, with zero representing no overlap. Whenever the most common pixel label pertained to GT background the associated IOU was set to zero, since no majority cell type has been identified.

#### Jensen-Shannon divergence

To evaluate how well the population of cells generated by the process of segmentation resemble the distribution of cell-areas of the GT we calculated the Jensen-Shannon divergence (JSD) as per R package *philentropy* (version 0.5.0) ^17^ that was used for the calculation:

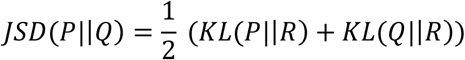

Where:

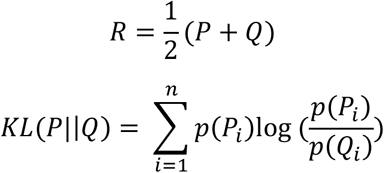

If the distributions are identical, the JSD is zero.

#### Root Mean Square Error

In the context of evaluating how well the two segmentation protocols can detect the abundance, both in terms of count percentage and area percentage, of different cell types, we calculated the root mean square error (RMSE) as follows:

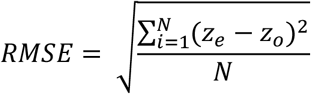

Where *N* is the number of cell types, *z*_*e*_ and *z*_*o*_ are the count percentage or the area percentage of GT and the product of segmentation respectively.

#### Diversity

To evaluate the difference in cell composition of a tissue induced by the two segmentation pipelines we borrowed the concept of diversity from ecology defined as follow:

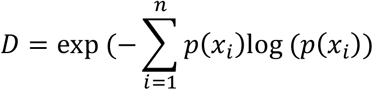

#### Importance

Accordingly to the R package Caret ^18^, used for the calculation, the importance of each parameter in a linear model is determined by the absolute value of the t-statistic.

## RESULTS

### Simulated Stress test

In order to evaluate the response of the two pipelines to a few common situations that might occur in histological staining such as the occurrence of regions with different cell density or the presence of cell and structures that heavily deviate from a generic rounded shape, we produced synthetic data that simulate the increase of a specific stressor.

In the first test (fig. 2) we arranged a series of images composed by same cell types (as presented in figure 1B) and identical proportions between them, but with increasing cell density (figure 2.A). We evaluated the reaction of the two segmentation processes quantifying the entropy and the IOU for each discovered cell and the Jensen-Shannon divergence of the cell area distribution of each prediction against the cell area distribution of the GT. In other terms, IOU and entropy are used to evaluate the goodness of prediction at the cell level while JS divergence is used to evaluate the prediction at the population level (fig 2B). Although partially correlated, gain of entropy, interpreted as loss of information^19^, and IOU contraction, as a more intuitive measure of observed-expected spatial coincidence, were both taken into account. Both pipelines presented very similar trends of gain of entropy (fig 1B upper left panel) and loss of IOU (fig 2B middle left panel), although the total effects, expressed as the sum of entropy and IOU values for the whole range of density in three independent determinations, was not significantly different (fig 2B right panel). Interestingly, the progressive change of JS divergence appears, on the contrary, quite different, with opposite effects around 50% of cell density. This trend can be explained as follows: at lower densities both pipelines discover nearly the same number of cells with the *reference* pipeline performing better in matching the correct extent (this is also corroborated by a slightly higher IOU, but also a marginally increased entropy). With the increase of cell density both pipelines start losing the ability to discover the correct number of cells, approximatively at the same rate, while the ability of covering the correct number of pixels seems to affect the *reference* pipeline more heavily than the *alternative* (fig 2B lower panel and supplementary fig 2.1). The overall outcome of this test demonstrated no significant differences between the two approaches, while suggesting a slightly better performance of *reference* over *alternative* at lower density, balanced by an opposite effect at higher density.

**Figure 2:**
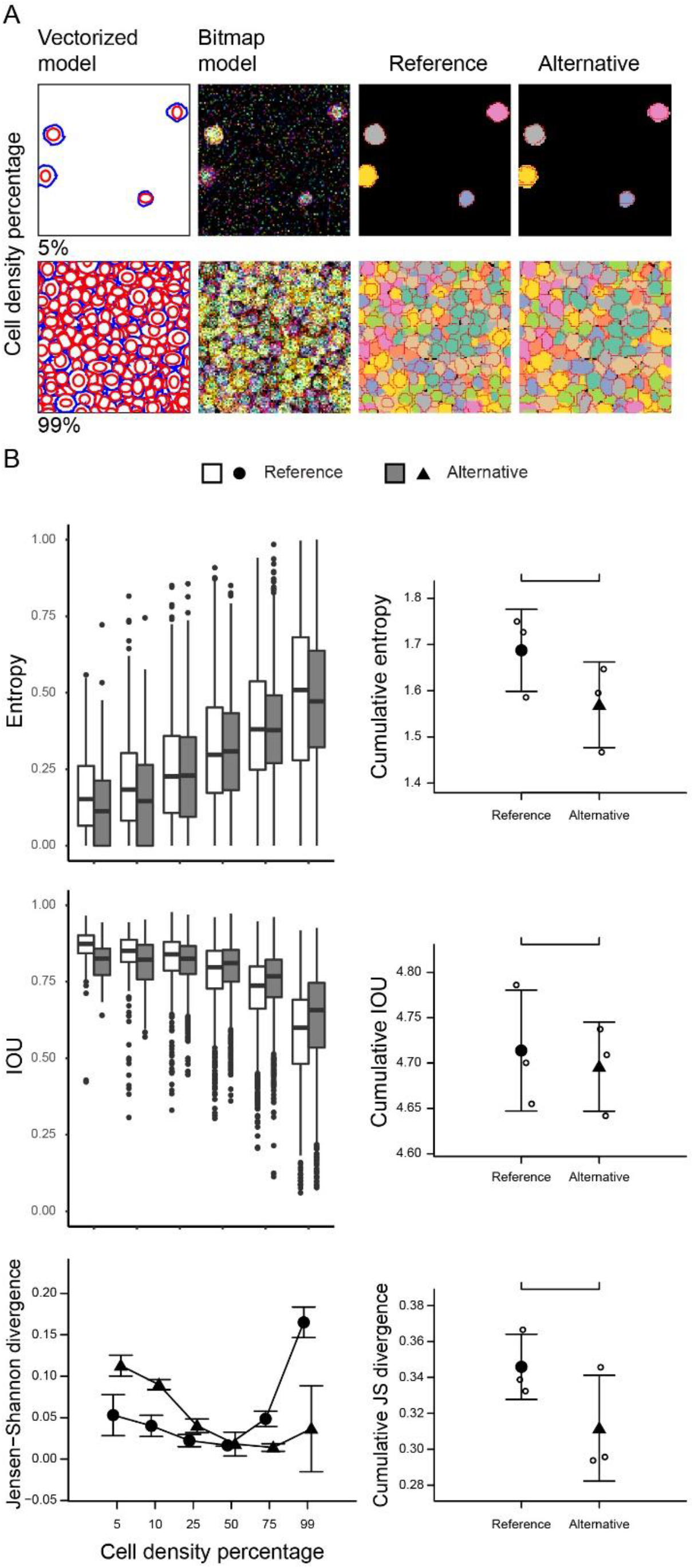
effect of cell density. Synthetic data has been constructed sequentially increasing density of cells from 5% to 99%-pixel occupancy of a 500 × 500 pixel image. The training process, for both pipelines, has been carried out on a portion of the image at 50% density. Representative images (100 × 100 pixels) of the lower and upper density range limits (A), from left to right each image represents the vectorized model, the overlay of the six simulated markers (cytoplasmatic common to all the cells (red), sharp plasma membrane marker (green), soft plasma membrane marker (blue), cytoplasmic marker (gray), cytoplasmic marker with two different mean intensity (cyan), and nuclear marker (magenta)), the GT overlayed with the *reference* segmentation, and the GT overlayed with the *alternative* segmentation. Quantification of entropy, IOU, and Jensen-Shannon divergence(B); for all the three metrics panels on the right shows the trend in one representative test, while panels on the left represents the cumulative value. Data are presented as mean±SD, n=3 independent tests and the statistics is a two-tailed, paired t-test codified as: “” p≥0.1,. “.” p<0.1, “*” p<0.05, “**” p<0.01, “***” p<0.001.

For the two subsequent tests, we decided that fixing the cell density at 75% would be a fair compromise considering the response of the two pipelines to the previous test and what would have been a realistic situation (considering the evaluation presented later in fig 6A and 7A).

Real cells could be considered to be mostly round in shape, especially immune cells, with some notable exceptions such as polygonal structures, for example in epithelial cells, or a dendritic profile for follicular dendritic cells or an elongated aspect for endothelial cells. To test the adaptability of the two protocols to a consistent and progressive deviation from an even sub-circular structure of cells we tested their predictive ability against a dataset of five images in which the compactness of cells, expressed as the ratio of the area of the actual polygon and the area of the associated convex hull vary from 0.97 (rounded cells) to 0.28 (dendritic-like cells, fig 3A). Similarly to the previous test, the trend change of both entropy gain and IOU loss for the two appear similar (fig 3B upper and middle left panel), with an overall significant better performance of *alternative* over *reference*. Again, the JS divergence revealed some more interesting results showing how, apart from the initial displacement of the two curves, the *reference* pipeline showed a steeper gain of divergence in comparison to *alternative* (fig 3B lower panel).

**Figure 3:**
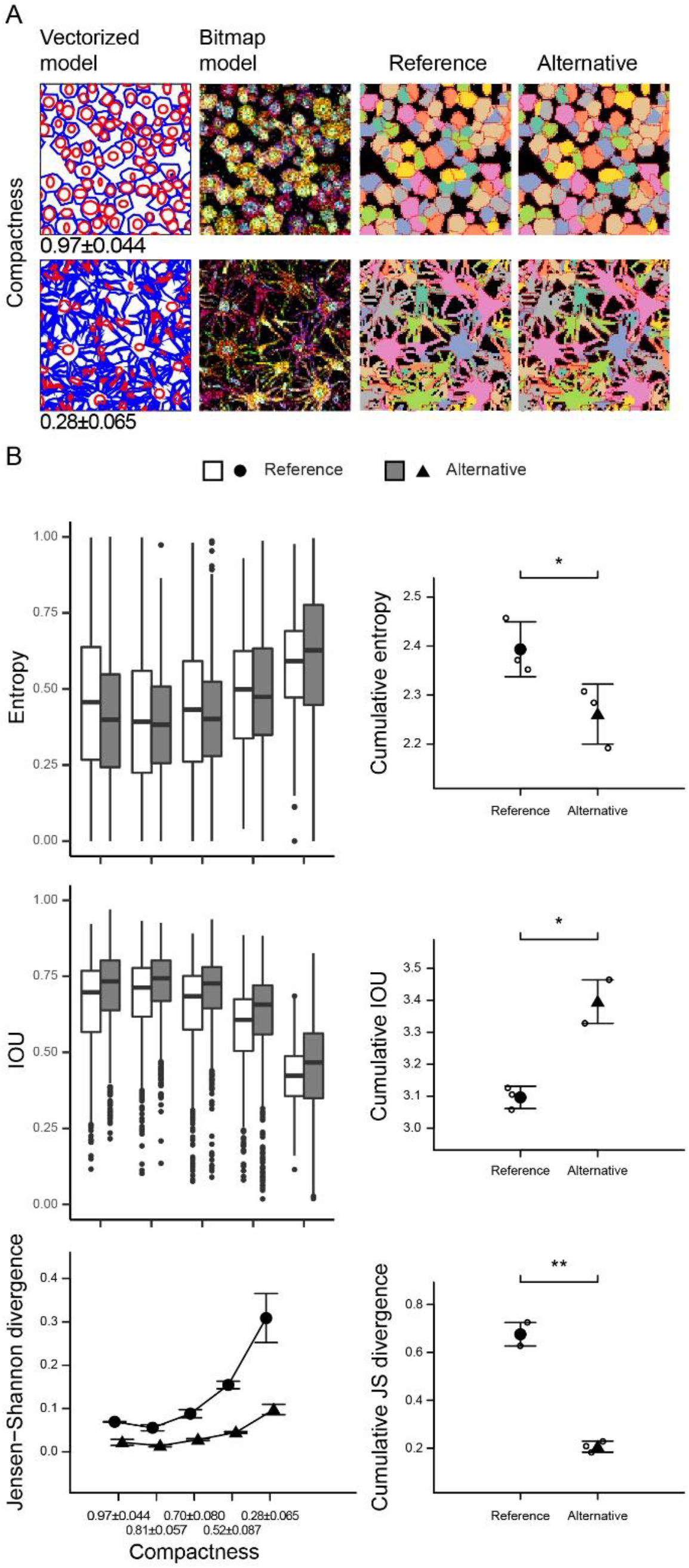
effect of geometrical complexity. Synthetic data has been constructed sequentially decreasing the compactness, expressed as the area-ratio of the polygon representing a cell and the associated convex hull, from 0.97±0.044 (round cells) to 0.28±0.065 (irregular cells with long dendritic protrusions). The cell density was kept fixed at 75% pixel occupancy of a 500 × 500 pixel image. The training process, for both pipelines, has been carried out on a portion of the image at the range point corresponding to 0.70 compactness. The training has been independently performed for the three repetitions. Refer to figure 1 legend for further details.

Another parameter that may vary between acquisitions is the quality of the images. In the third test we constructed a dataset similar to that used in the first test at 75% of cell density and progressively deteriorate the signal by adding random noise. Differently from the previous test, where the stressor was supposed to affect mainly the process of segmentation, this test is expected to mainly affect the classification step carried out before segmentation. The overall outcome (fig 3B left panel) indicated a significant better performance of *alternative* over *reference* in at least two parameters out of three, whereas the divergence analysis showed an extremely different behavior of the two pipelines. *Reference* demonstrated to be extremely stable throughout the test while *alternative* showed to be sensibly affected after the amount of noise increased beyond 20%. Although without an empirical proof, we would argue the difference could be related to the fact the two models would be differently sensitive to noise because of the complexity of *alternative*, that is composed by several different classes, in comparison to *reference* that is constituted by two simple independent binary systems (nuclear-staining/ background and cytoplasmic-staining/background as independent models).

### Simulated discrimination test

A common task in an imaging experiment is the discrimination of different cell subsets in a complex mixture. To evaluate the performance of the two pipelines we arranged a test where different cell types, with complex phenotypes such as in the definition of T-cell subsets (naïve, effector, memory, etc.), were interspersed in a relatively homogeneous agglomerate of cells or simulated tissue. Target cells were designed to have mainly a random shape, while tissue cells were constructed as slightly bigger polygons to simulate a parenchymal structure infiltrated by immune cells, for example an epithelial tissue (as presented later in this paper) or an organ with a highly repetitive infrastructure such as liver. Obviously, these simulations do not take into account the organisation of higher-level structure, such as the different layers of an epithelial tissue or the organisation of a follicle and mantel zone as found in lymph node (also presented later in this manuscript), that would have required a much more complex algorithm which would have exceeded the scope of these tests.

From a first empirical comparison of the reconstruction obtained from the *reference* pipeline (fig 5 A left panel) and *alternative* pipeline (fig 5A middle panel) with GT (fig 5A right panel) it appears quite evident *alternative* resemble the overall ‘tissue-structure’ more closely than *reference*. It was noticed that both pipelines showed a tendency to ‘inflate’ the area of the target cells to the detriment of the surrounding ‘parenchyma’; this is also confirmed inspecting the boxplot (fig 5B, first and second row) representing the total area per cell type, in which all the areas (a part from cell type D.2) are overestimated by both pipelines, while the area covered by cell type E is always underestimated. The opposite effect can be appreciated when we compare cell counts (fig 5B, third and fourth row). As per the overall effect, evaluated in terms of the root mean squared error (RMSE) and the JS divergence of the distribution of area extents of both pipelines given the same distribution of GT, *alternative* performed better than *reference* (fig 5C).

**Figure 4:**
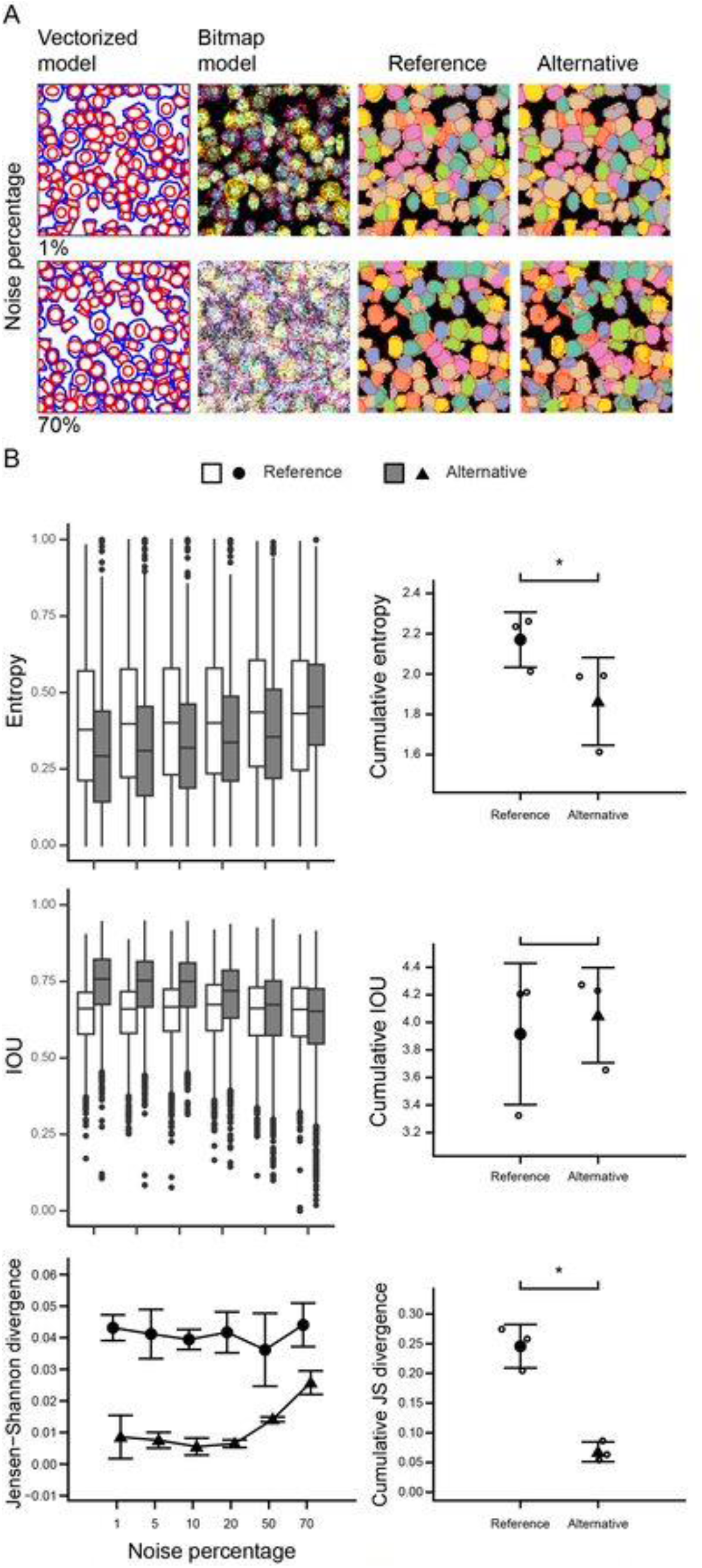
effect of scattered noise. Synthetic data has been constructed sequentially increasing the number of pixels modified by the addition of a noise signal to all the markers, forming the layer stack, as per a random amount selected within the normal distribution having as mean and standard deviation the same mean and standard deviation of the entire layer to which it was applied. The cell density was kept fixed at 75% pixel occupancy of a 500 × 500 pixel image. The training process, for both pipelines, has been carried out on a portion of each image and combined in a single training set. The training has been independently performed for the three repetitions. Refer to figure 1 legend for further details.

**Figure 5:**
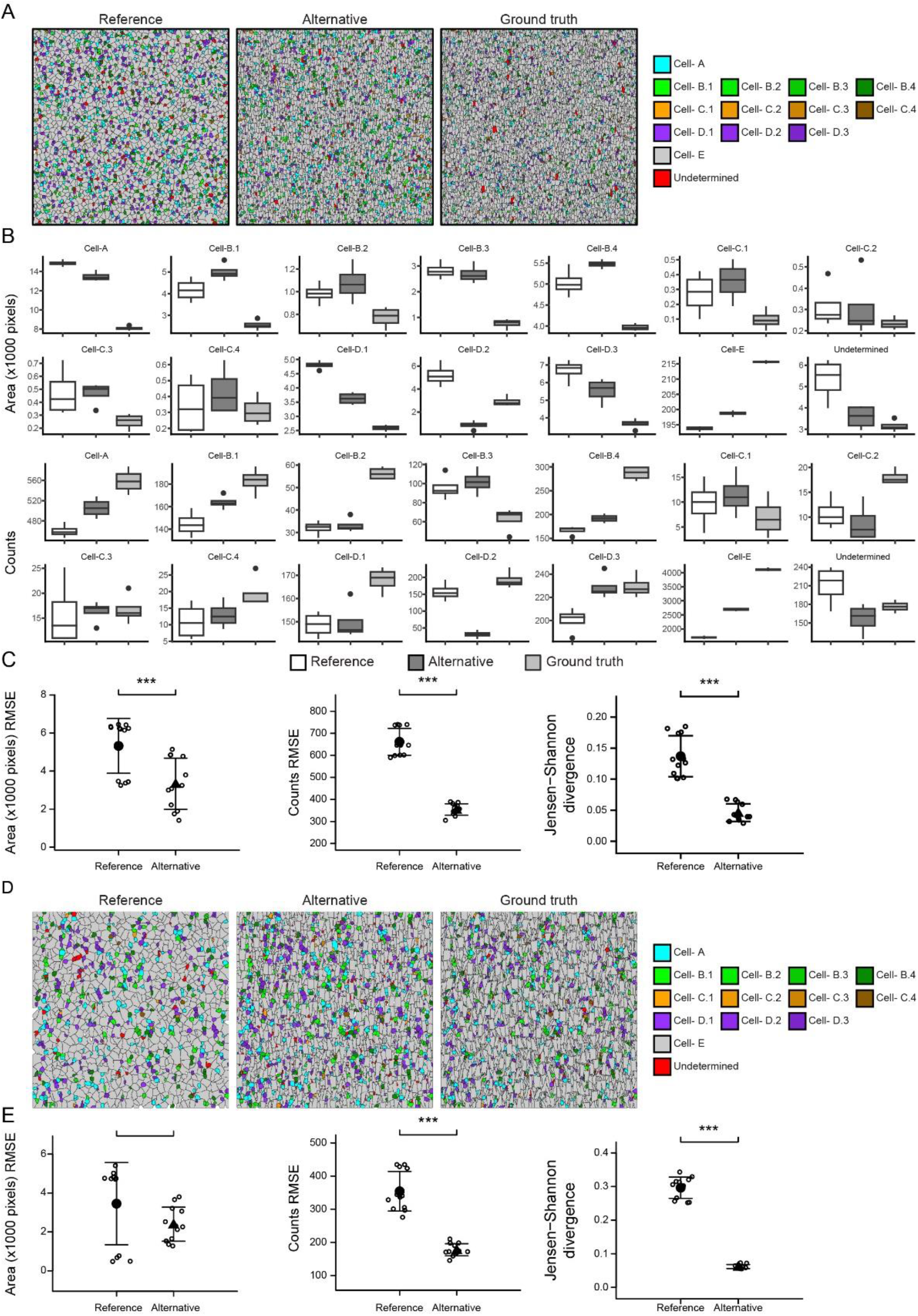
efficiency of cell type identification in a complex mixture. Two series of 12 images (500 × 500 pixels) have been constructed as a mixture of 13 cell types with different cell markers, simulating plasma membrane, cytoplasmic and nuclear distribution with either a single mean value or a mixture of high and low expressing cells. The two series have similar characteristics in terms of cell composition, and both have been set to have a cell density of 98% but have different median cell size, with smaller cells (A, B and C) and larger cells (D and E). Representative image of segmentation results are shown in (A) and (D) for small and large cells respectively, from left to right: *reference, alternative* and GT. Each series of 12 replicates has been divided in three subsets for which the training process has been carried out independently on a portion of one image. Boxplots (B) representing the median, inter-quartile range, min-max range, and outliers for one representative set show the difference in area (first and second row) and counts (third and forth row) within each cell types for the *reference* (white box) and *alternative* (dark gray box) estimations as compared to the expected value (pale gray box). Evaluation of the root mean squared error (RMSE) for the *reference* (filled circle) and *alternative* (filled triangle) for the area and count estimation, and the Jensen-Shannon divergence against the GT area distribution for respectively the *reference* (filled circle) and *alternative* (filled triangle) segmentation. Data are presented as mean±SD, n=12. The statistics is a two-tailed, paired t-test codified as: “” p≥0.1,. “.” p<0.1, “*” p<0.05, “**” p<0.01, “***” p<0.001.

A second dataset has been created similarly but with proportionally larger cells with the idea that this factor would have influenced the performance of segmentation. As depicted in fig 5D the geometrical distortion noticed in particular for *reference* still affects the prediction, none the less the associated estimation errors decreased for both area and counts at the point that the difference in performance between the two pipelines, although still significant in terms of counts, disappear when we consider the areas. The difference in JS divergence remained significant and interestingly showed a slight increase, mainly affecting *reference*.

### Case study 1: staining of FFPE histology from 3 follicular lymphoma patients

Follicular lymphoma (FL) is a haematological malignancy of the lymphoid tissue characterized by germinal centre (GC) B-cell differentiation and it is characterized by a high degree of heterogeneity comprising several morphological variants^20^. Lymph nodes are complex anatomical structure where the lymphoid lobule constitutes the basic anatomical and functional unit, compartmentalized as cortex, paracortex and medulla. Further compartmentalization of each lobule generates distinct areas for B and T lymphocytes to interact with their specific antigen presenting cells (APC): follicular dendritic cells (FDC) and dendritic cells (DC) respectively^21^.

Lymph nodes present an interesting example of a cell crowded tissue (fig 1C upper panel and supplementary fig 6.1 upper row) in which even the boundaries between cell nuclei are fuzzy. We analysed three ROI from three different patients, sampling a similar portion of each image for both pipelines. As per the previous tests the training strategy for the *reference* pipeline was to train separately one random forest model for the contrast DNA/ background and one for a mixture of the remaining markers versus background. On the contrary, for the *alternative* pipeline random forest was trained on the following classes: CD4^+^, CD8^+^, CD20^+^, CD21^hi^, CD68^+^, CD16^+^, CD31^+^, ACTA2^+^, and background.

Segmentation resulted in a significantly higher total area classified by *alternative* as compared to *reference* (fig 6A). In order to understand the marker distribution produced by the two segmentation algorithms, we plotted all pairwise scatterplots for the two protocols (fig 6B and supplementary fig 6.2). The most striking difference in the staining patterns was that *alternative* more easily identified discrete subsets of positive cells (this statement is generally valid for all the markers but CD20 probably because of the diffuse staining this marker produced, indicating the necessity of adjustments in the staining conditions for this marker) as compared to *reference*. This effect offers, in the process of multivariate classification, the advantage of making the selection of threshold values easier. Two possible factors can contribute to the appearance of discrete population in the *alternative* segmentation: first *alternative* segmentation is conducted separately on subsets of pixels accordingly to the first round of classification, this increases the chance of isolating more ‘pure’ areas; second, the constrain on the nuclear marker imposed by the *reference* segmentation can result in this being too stringent in certain situations, although it appears to be relevant preferentially with certain markers such as CD21 and CD68 (supplementary figure 6.3), for which a nucleus can possibly not be evident due to the irregular shape of this dendritic-like cells.

**Figure6:**
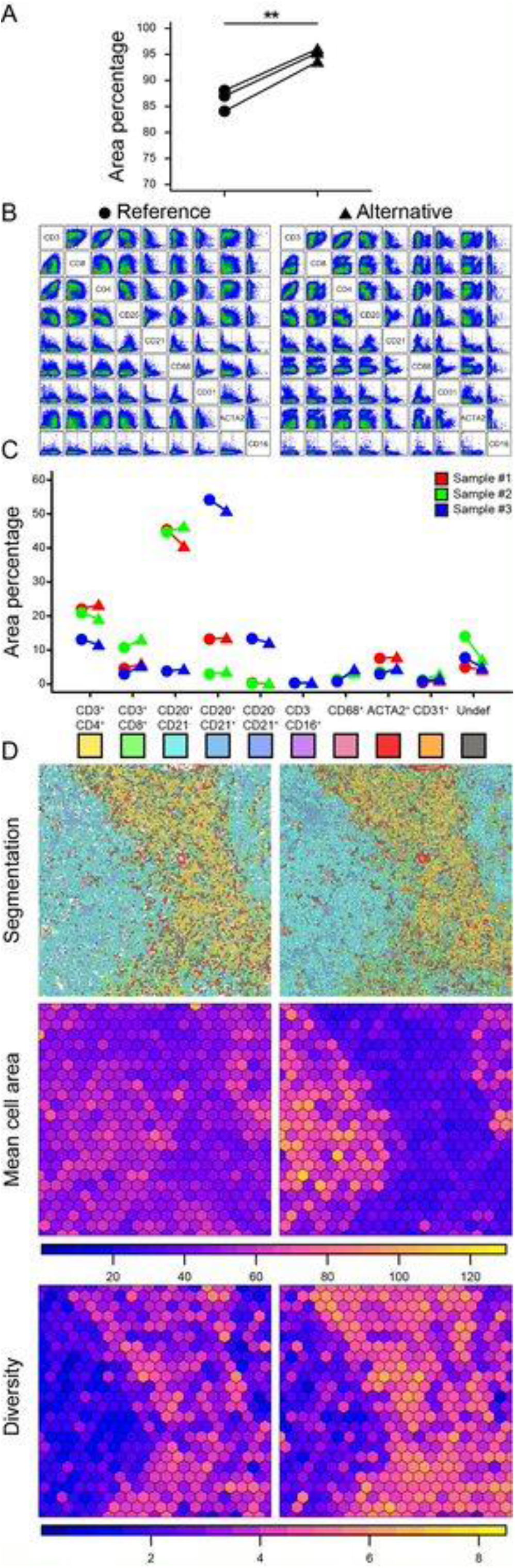
case study 1, segmentation of follicular lymphoma FFPE histology. Total segmented area for *reference* (filled circle) and *alternative* (filled triangle). N=3 independent patients, statistics is a two-tailed, paired t-test codified as: “” p≥0.1,. “.” p<0.1, “*” p<0.05, “**” p<0.01, “***” p<0.001 (A). Representative bivariate scatter plot of the mean intensity for each marker in sample #2 (B). Dashed vertical and horizontal lines represent the threshold values used in the definition of cell subsets. Area percentage on total segmented area for each of the cell subset identified in three different samples (C). None of the subset is significantly different in a multiple-pairwise paired t-test with Bonferroni correction. Segmentation results for sample #2 obtained with the *reference* pipeline (left, upper panel) and *alternative* pipeline (right, upper panel), mean cell area (middle panel) and diversity (lower panel, D)

Although no significant difference between the two segmentations was noticed in the relative amount of area covered by each cell class (fig. 6C), we observed that the *alternative* protocol was biased towards the detection of higher local diversity (fig. 6D lower panel) or conversely *reference* understated diversity. The particular formulation that we used to evaluate diversity, corresponding to what in the field of ecology is known as Hill numbers formulation for q=1 can be accounted as the effective number of species in a group or collection ^22^. Since we know from the previous tests that the two segmentations algorithm can produce very different area distributions, in order to evaluate whether a gain in diversity may or may not be a direct effect of smaller segments, we projected the mean cell area per tile as well. As presented in figure 6D middle panel, *alternative* segmentation produced a more widespread range of areas in sample #2, while both sample 1 and 3 were more biased towards smaller areas in *alternative* segmentation as compared to *reference* (supplementary figure 6.5). The discrepancy in area extent, in absence of a ground truth reference, could be seen either as over-segmentation or under-segmentation, although both phenomena, if producing a homologous transformation in which, for example, the over-segmentation products belong to the same class of the bigger parental class, would not directly traduce in a gain of diversity, since diversity, in this formulation, is only governed by the relative proportions of each class. As a matter of fact, if we analyse the local presence of a class, counting the number of tiles where one type of cell appears within a specific tile in one of the two protocols but not in the other (supplementary figure 6.5), we noticed, accordingly to the observed higher diversity, that the *alternative* protocol is able to constantly catch more unique types with a conserved peak on segments classified as CD68^+^, with a prevalence of almost 50% of tiles that contain this type of segment uniquely in the *alternative* segmentation. As pointed out before the alleged increased sensitivity of the *alternative* segmentation algorithm might be connected to the weaker constrains it has to nuclear staining as segmentation initiator (supplementary fig. 6.3).

### Case study 2: staining of FFPE histology from head and neck cancer patients

Head and neck squamous cell carcinoma (HNSCC) is a heterogeneous pathology which pathogenesis, in most cases, take place within large pre-neoplastic patches of mucosal epithelium^23^. HNSCC can present both high and low levels of immune cell infiltration and immunological activation which showed correlation to various factors such as smoking history (immunosuppression and lower levels of cytotoxicity) human papilloma virus (HPV) positivity (positive correlation)^24^.

We arranged a panel of markers that would broadly cover the immunological infiltrate or ‘immune contexture’ of a tumour and the possible location of tertiary lymphoid structure ^25,26^. We tested this panel on 3 or 4 distinct ROIs from two consecutive sections of two different HNSCC patients at early stage of tumour progression, for a total of 14 samples.

As in the previous case study, *alternative* protocol was able to detect and classify a larger portion of the image as compared to *reference* protocol (fig 7A). In this occasion, instead of a rigid classification of cells based on hard thresholds we applied a consolidated analysis (at list in flowcytometry and liquid mass cytometry applications) based on the use of the clustering tool FlowSom ^27^. Data has been transformed with a modulus transformation^28^:

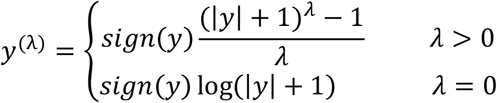

**Figure7:**
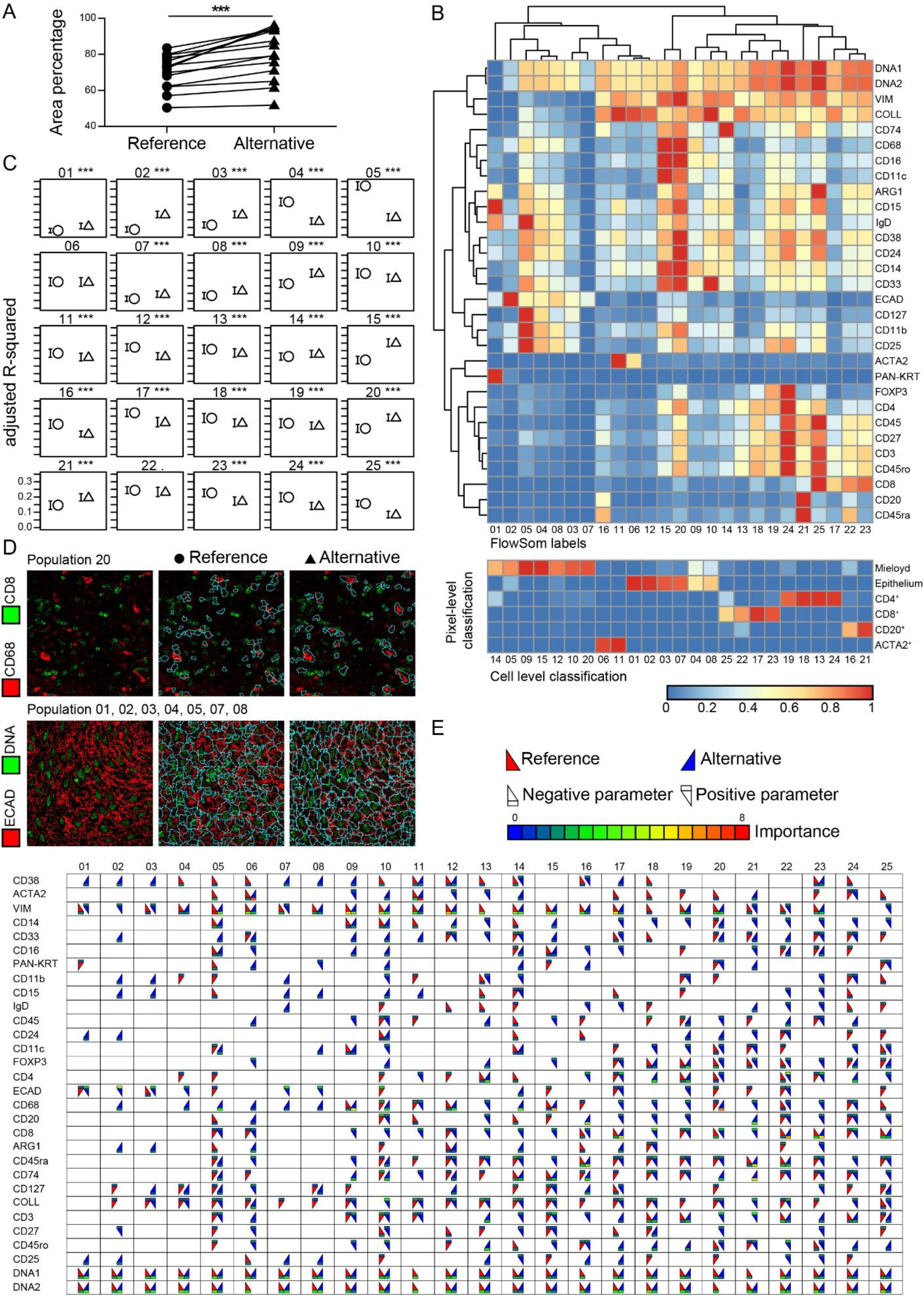
case study 2, segmentation of head and neck FFPE histology. Total segmented area for *reference* (filled circle) and *alternative* (filled triangle). N=14 independent ROI from two different patients, statistics is a two-tailed, paired t-test codified as: “” p≥0.1, “.” p<0.1, “*” p<0.05, “**” p<0.01, “***” p<0.001 (A). Both the pipelines have been trained on the same portions of three different ROI and analyzed separately. The segmented cells from the two pipelines have been collated and clustered using the R package FlowSOM. Heatmap representing the scaled mean fluorescence intensity (MFI) for each marker (rows) in each of the 25 SOM clusters (columns, B upper panel). Heatmap representing the relationship between the biased classification used for the *alternative* training (pixel-level classification, rows) and the FlowSOM clustering (cell-level classification, columns, B lower panel). Comparison of the adjusted R-squared value for the linear regression analysis in which the position of each pixel within a polygon is used as the response variable and each marker of the staining panel as explanatory variables. N= 100 bootstrap repetitions on a subset of n=100 cells sampled with replacement. statistics is a two-tailed, unpaired t-test with Bonferroni correction and codified as: “” p≥0.1, “.” p<0.1, “*” p<0.05, “**” p<0.01, “***” p<0.001. (C). Representative histological images showing the staining for CD68 (red) and CD8 (green, upper row), and E-cadherin (red) and DNA (green, lower row), without segmentation (left column) or overlayed by the *reference* segmentation (central column) and *alternative* segmentation (right column) for selected FlowSOM subsets (D). Pictogram synthesizing the direction (upright triangles: negative parameter, downward triangles: positive parameter) and importance of each parameter of the linear model. Only the parameters statistically significant (p<0.05) are represented (E).

Using for this data set λ=0 that traduces in simple logarithmic transformation of the data shifted by one unit. After data sets from the two segmentation analyses have been collated, FlowSOM algorithm has been instructed to arrange a self-organizing map (SOM) of 5 by 5 clusters. Analysis result is presented as heatmap (fig. 7B, upper panel) of the mean intensity values of each marker rescaled to the range 0 to 1. The 25 resulting clusters can be roughly interpreted on the basis of the column dendrogram as the left upper branching pertain to non-immune cells while the right one aggregate mostly immune cell types, where we can notice a progressive change of cell types from CD8^+^ lymphocytes on the extreme right of the plot, progressively through CD20^+^, CD4^+^ lymphocytes and myeloid cells moving towards the middle.

In order to check what would be the relationship between the pixel-level classification, obtained from the pre-segmentation random forest analysis, and the FlowSOM clusterization (this was possible only for the *alternative* protocol since the *reference* random forest was constructed using a simple dichotomic definition), we enumerated the cells for each of the combination between the classes used for the *alternative* random forest training (pixel-level classification) and the 25 clusters obtained from the FlowSOM analysis (cell-level classification) and organized the results as a heatmap with the cell counts rescaled by columns to the range 0 to 1 (that can be interpreted as the probability of the pixel-level classification given the cell-level classification, Fig 7B lower panel). The concordance between the two classification was generally high (red tiles), with few FlowSOM clusters presenting a more spread distribution across rows. In particular, we noticed that some level of uncertainty exists between myeloid cells and epithelial cells, and between CD8^+^ lymphocytes and both CD4^+^ and CD20^+^ lymphocytes.

To interpret the FlowSOM clusters from a different perspective, we analysed the linear relationship that might exist between the staining pattern of each marker with the position of each pixel inside the cell it belongs to. Pixel position has been calculated through a process of progressive erosion of each cell in which the outer pixels assume higher values and the most internal are zero. Since the two segmentations gave similar pixel-position distribution (supplementary fig. 7.1) but with a slightly broader range in the *reference* protocol, and in order to guarantee a population of pixels large enough to sample, we limited the regression analysis to the position range between 0 and 7. Each cluster have been analysed separately bootstrapping 100 times each linear regression conducted on a sub sample of 100 pixels for each of the eight possible positions. The comparison of the R-squared, represented as mean and standard deviation of the bootstrap, is presented in figure 7C and the synthetic representation of the trend of significant parameter in figure 7E. Only parameters that were significantly different from zero (p-value<0.05) were included in the pictogram. Negative parameters (markers that are brighter towards the centre of a cell) are represented by a triangle pointing upward, while positive parameters (markers brighter towards the edge of a cell) are identified by upside-down triangles. The general picture present markers that are either more important for one segmentation protocol over the other, or concordant or antithetical. As expected, the DNA-marker is important and concordant for almost all cell classes, possibly because of the unambiguous and good quality nuclear staining is. We focused on some peculiar classes that presented controversial results. The FlowSOM cluster number 20 that would be interpreted as a phenotype strictly correlated to a myeloid profile, because of the high positivity for the main monocyte/ macrophage markers such as CD68, CD33, and CD14, with some positivity for T-lymphocytes (fig. 7B), presented a significant negative correlation to the above-mentioned myeloid markers in the *alternative* segmentation while presented an antithetical positive correlation in the *reference* segmentation. From a visual inspection of a representative image to which we overlayed both segmentation (fig 7D upper panel), appears clear how this cluster probably aggregate different cell types identified by the two segmentation protocols, with in particular the *alternative* protocol surrounding more precisely cells with high CD68 expression (red) and excluding neighbouring CD8^+^ T-cells (green), while *reference* segmentation identified nearby regions more loosely bounded to the mentioned markers limits, possibly obeying more closely to the DNA staining. Another interesting case is the group composed by the FlowSOM clusters 01, 02, 03, 04, 05, 07, 08, that pertain specifically to the tumour/ epithelial set with some leakage towards a myeloid phenotype. All these clusters but 05 presents a significant positive correlation of e-cadherin (ECAD) in the *alternative* segmentation, while only cluster 01 present the same correlation, although showing a lower importance of this parameter, in the *reference* segmentation. From a similar visual inspection (fig 7D) it appears that the *alternative* segmentation can detect cell edges, in this situation, more accurately as compared to the *reference*.

## DISCUSSION

In recent years the explosion of omics, high throughput and high-content technologies applied to life science is posing new challenges in different applications including those exquisitely based on imaging, bringing to them the so called ‘curse of dimensionality’, that is the exponentially growing need for data as the result of additional data dimensions in order to avoid unreliable data distribution and sparseness^3^. Cell segmentation becomes an essential step in imaging analysis whenever the analysis itself is meant to establish relationships, either merely quantitative or topological, of different features that populate a certain space. The number of tools to accomplish image segmentation is constantly increasing with notable example such as CellProfiler ^8^ (used as the gold standard and *reference* tool in this article) and others ^29^. With this paper we are presenting and testing against a consolidated method a novel pipeline that addresses two main issues: first the lack of a package dedicated to cell segmentation within R, a scripting environment constantly increasing in popularity among life scientist, in particular for cytometry applications, although package such as Cytomapper^30^ have been recently released this still rely on the use of foreign tools such as CellProfiler and ilastik (components of the aforementioned gold standard pipeline) for classification and segmentation, and a watershed algorithm is implemented in the package EBImage^31^. Second, we experienced some difficulties in applying more complex classification strategies that involve the identification and use of separate cell classes neighboring a communal space.

The novel *alternative* pipeline and the *reference* pipeline against which we benchmarked our solution, rely on very similar principles: an image, consistent of a stack of data arrays, or a portion of it, is first manually annotated, in order to create a training data set. A random forest algorithm is then used to predict and apply the classification to new unknown data. The new image created by the classification processes is then segmented. On top of communal framework a few substantial differences take place. While *reference* uses the classification process to create a probability map of likelihood of association of each pixel to a specific category as opposed to background, and this probability is used directly for segmentation, *alternative* creates the same map on a multi-class definition and uses the probability to create a pre-segmentation space in which each pixel is segregated to specific region to which they belong to. A second random forest, trained for regression instead of classification, that use the position of each pixel as response variable and each of the markers (native or calculated) as explanatory variables, is used to create one specific map of predicted positions for each of the categories defined in the classification step. This reinterpretation of the classification-segmentation approach permits to overload or bias the actual imaging data with the pre-knowledge or interpretation of an expert eye, such as that of a pathologist, and could facilitate the subsequent segmentation with a broader pre-segmentation step.

The use of a calculated dimension such as the pixel position, determined by the random forest regression based on all the available markers, could offer some advantages over the use of a crude single marker or single-class classification probability. As suggested by both synthetic (fig 5) and empirical data (fig 6), the adherence of the segmentation to the shape of each single cells can be improved by a biased approach. Using the nuclear staining as the trigger or seed initiator of segmentation is a common practice, although some drawbacks are implicit, such as the fact that in a tissue section of 4-7 µm thickness a proportion of cells will appear devoid of nucleus. Our *alternative* approach that does not rely strictly on the nuclear staining, although this remains one of the markers being constantly significant as demonstrated in fig. 7E, demonstrated to be able to explain and segment a statistically significant larger area as compared to *reference* in both the case studies we presented in this paper.

We demonstrated that the capacity to integrate into a classification/ segmentation model an amount of valuable information, such as the interpretation of different cell lineages given by an expert eye, contributed to a better segmentation. This poses the basis for a new task i.e., to model pathologist’s notions beyond the single cell level towards the physiological and pathological tissue architecture.

## Acknowledgments

This work was supported by Cancer Research UK funding support to King’s College London – UCL Comprehensive Cancer Imaging Centre (CR-UK & EPSRC), Cancer Research UK King’s Health Partners Centre at King’s College London, and Cancer Research UK UCL Centre; as well as CRUK City of London Centre.

LD is supported by Multidisciplinary Project (Award number 21840) and EU IMI2 IMMUCAN (Grant agreement number 821558)

PRB is supported by the Cancer Research UK King’s Health Partners Centre at King’s College London, and Cancer Research UK UCL Centre; University College London.

KN is supported by Cancer Research UK Clinical Training Fellowships (Award number 176885).

PP is part funded by MRC Clinical Academic Research Partnership (Award number MR/T005106/1)

RM is supported by Cancer Research UK King’s Health Partners Centre at King’s College London, King’s College London – UCL Comprehensive Cancer Imaging Centre (CR-UK & EPSRC), and CRUK NCITA: National Cancer Imaging Translational Accelerator ((Award number 28682).

JD is supported by a KCL-China Scholarship Council PhD studentship.

We wish to thank Joson Mathew for testing RUNIMC and being one of the first users.

## List of supplementary materials

**Supplementary figure 2.1:**
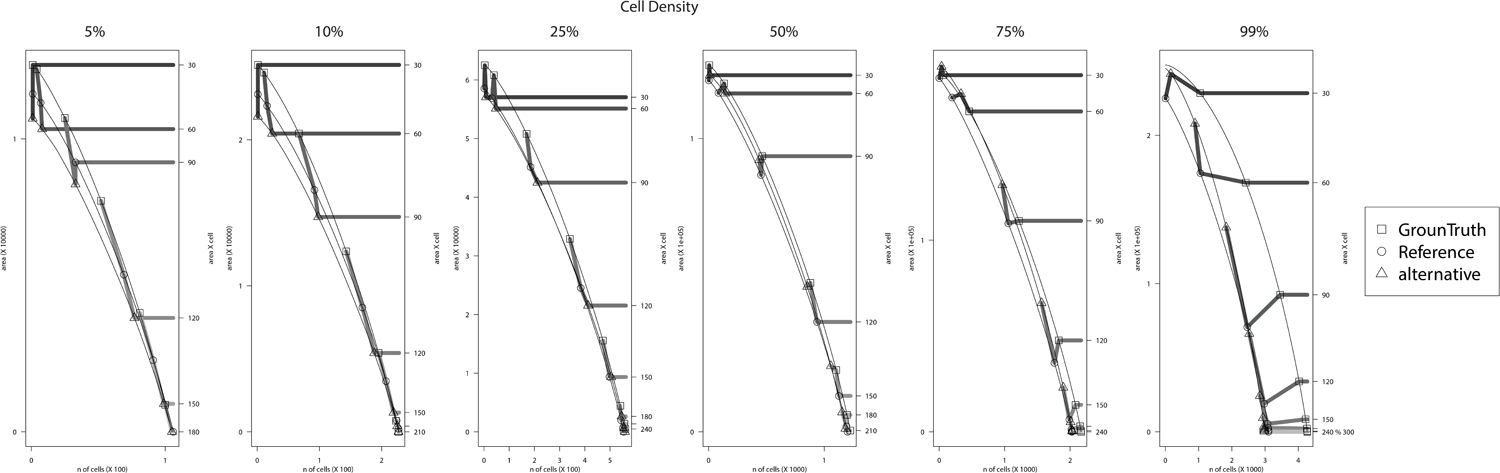
support to figure 2.B lower panel. Distribution of cell area for each of the six cell density levels.

**Supplementary figure 6.1:**
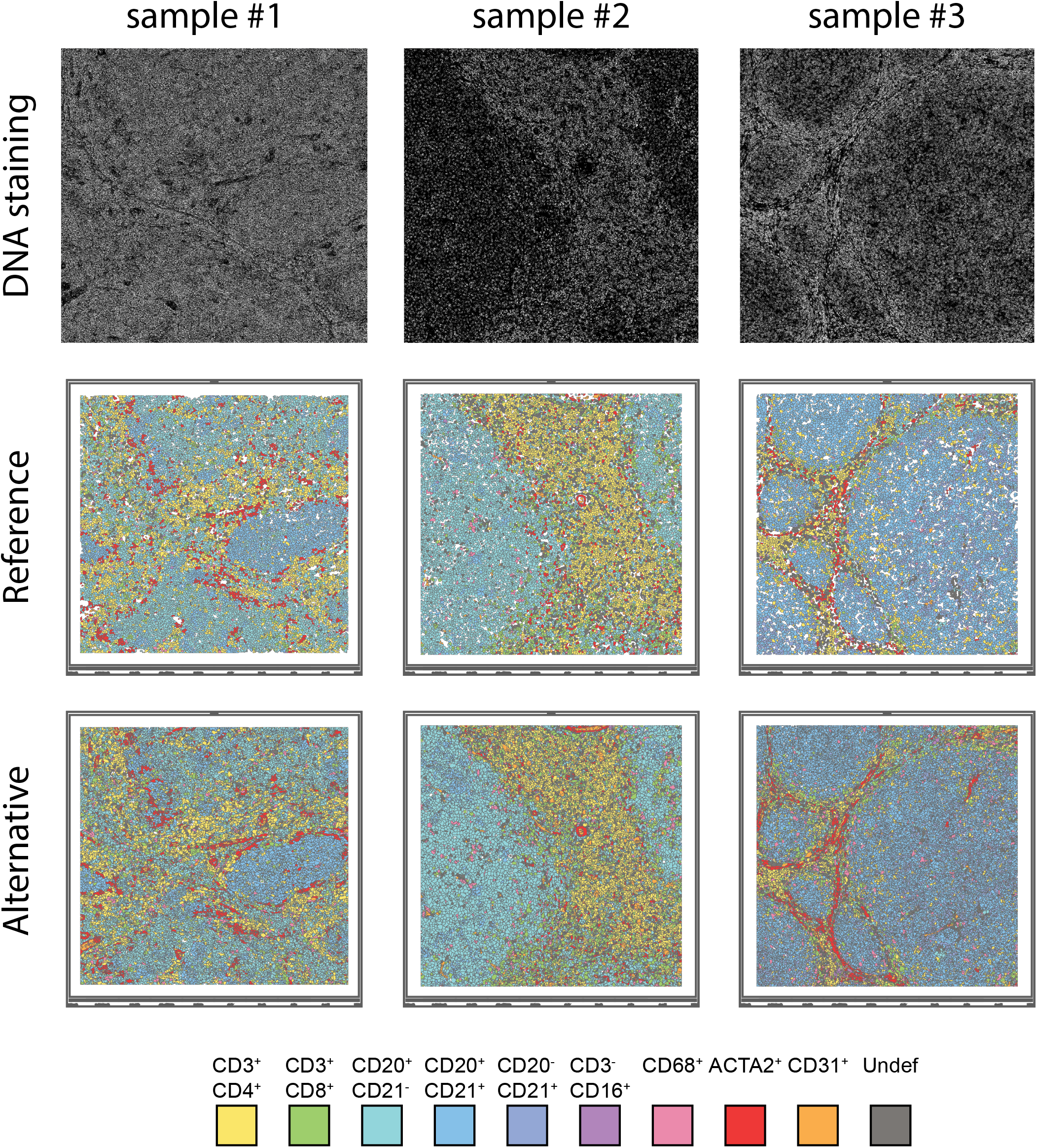
support to figure 6. Segmentation results for the *reference* and the *alternative* pipeline for the three lymphoma patients.

**Supplementary figure 6.2:**
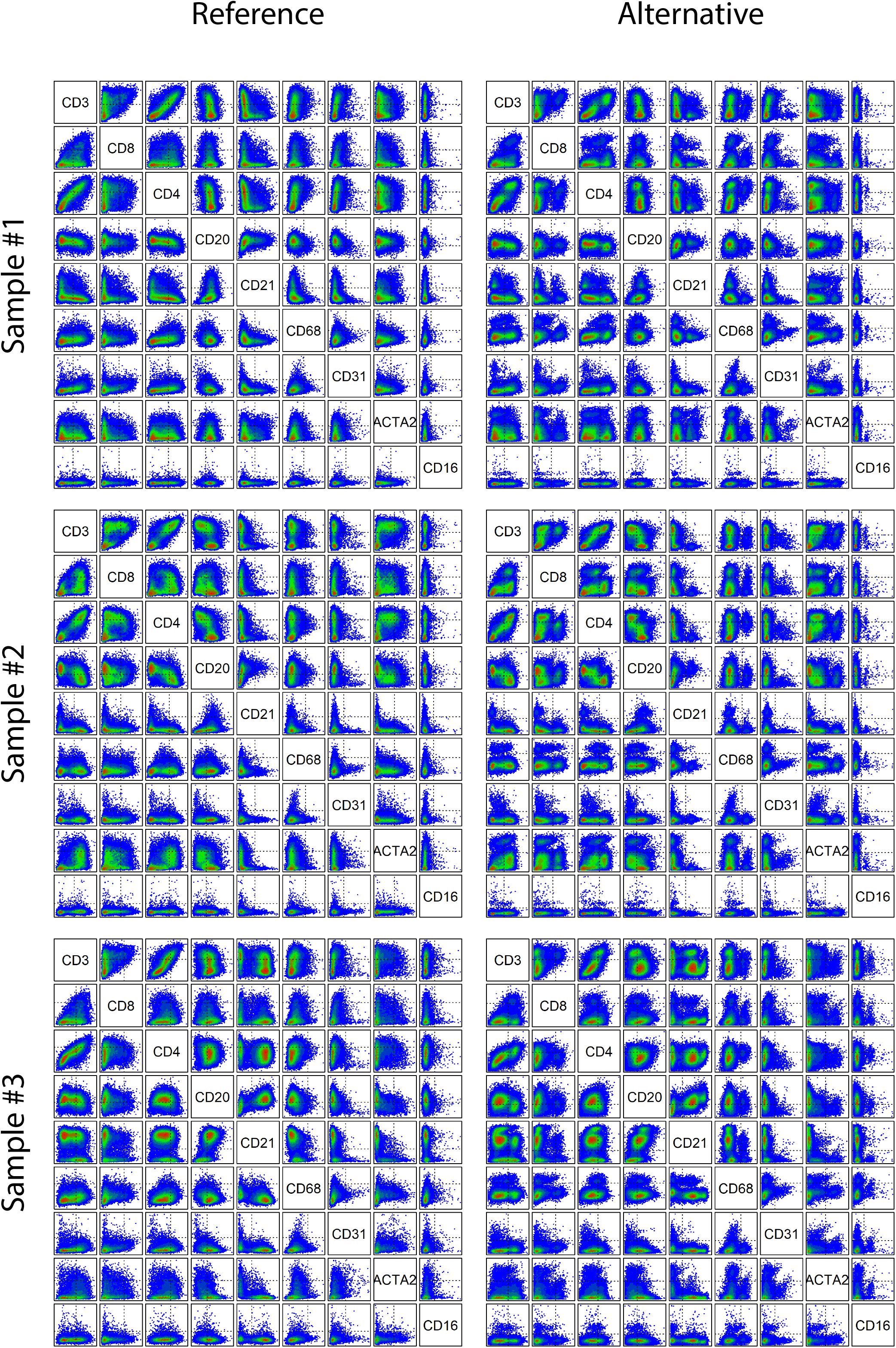
support to figure 6B. Scatterplots of marker combinations for the three lymphoma patients.

**Supplementary figure 6.3:**
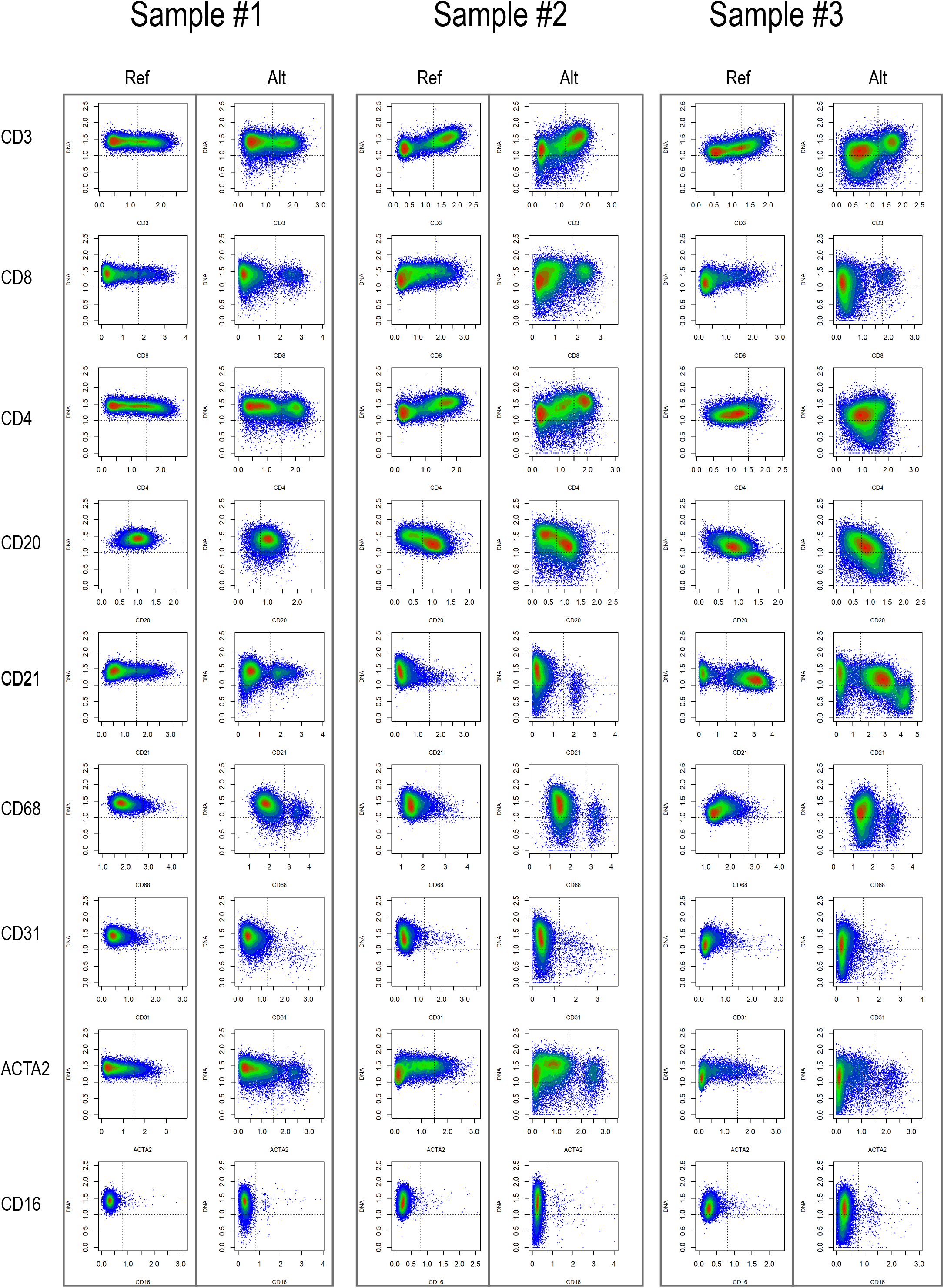
support to figure 6B. Scatterplots of each marker versus the DNA staining for the three lymphoma patients.

**Supplementary figure 6.4:**
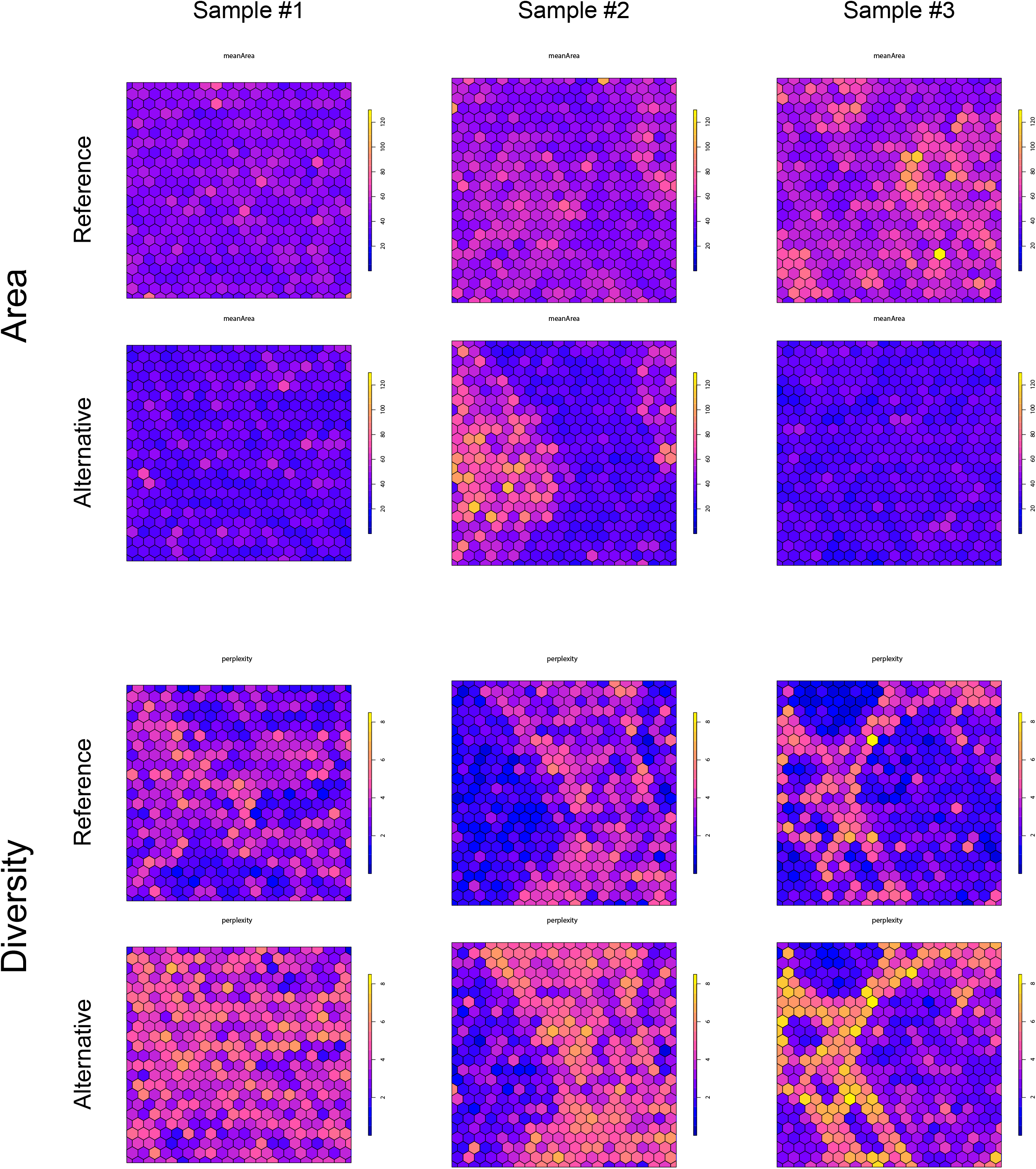
support to figure 6D middle and lower panel. Mean cell area and diversity estimation for the three lymphoma patients.

**Supplementary figure 6.5:**
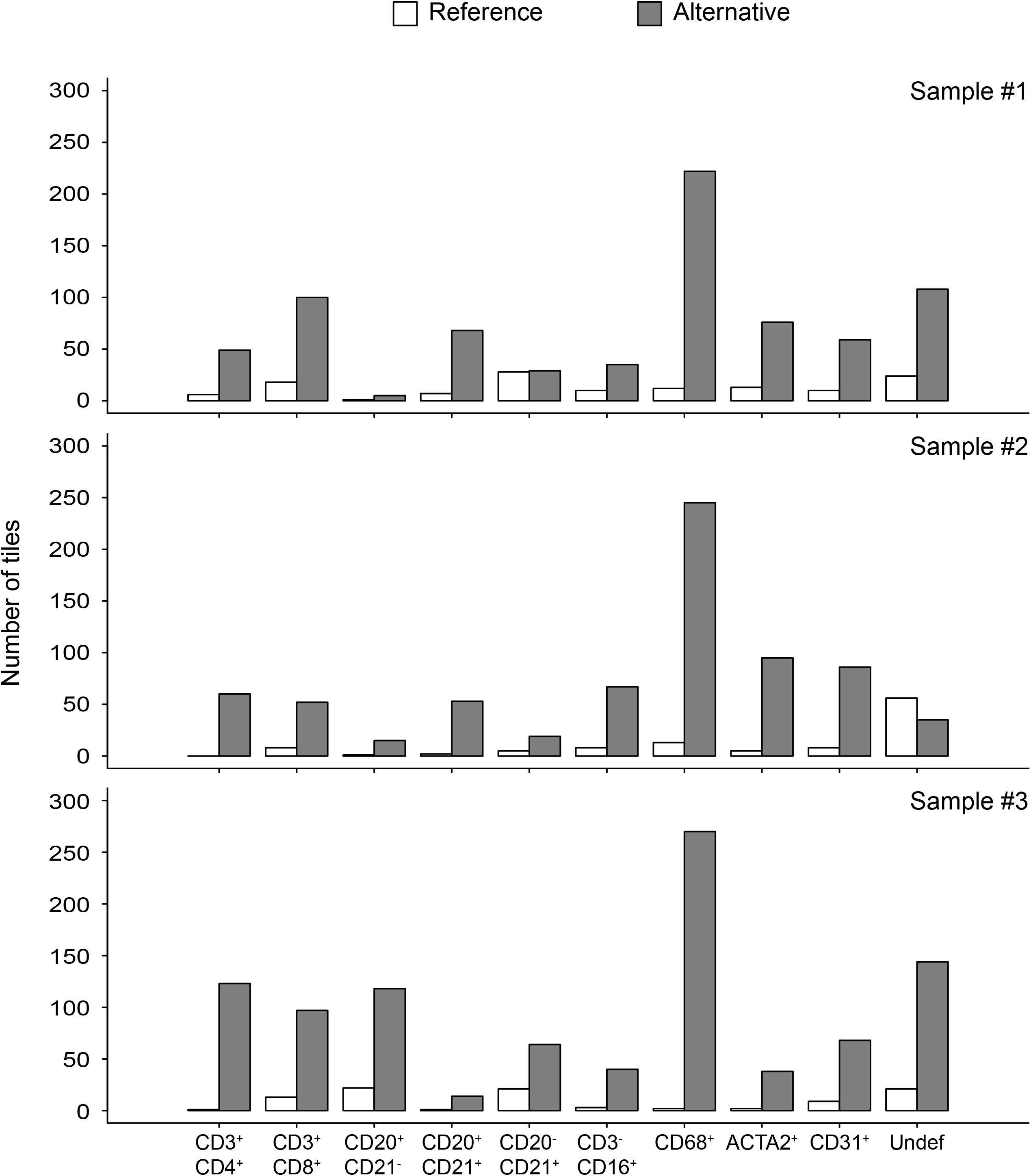
support to figure 6D lower panel. Cell type per honeycomb cell appearing exclusively in either *reference* or *alternative* segmentation

**Supplementary figure 7.1:**
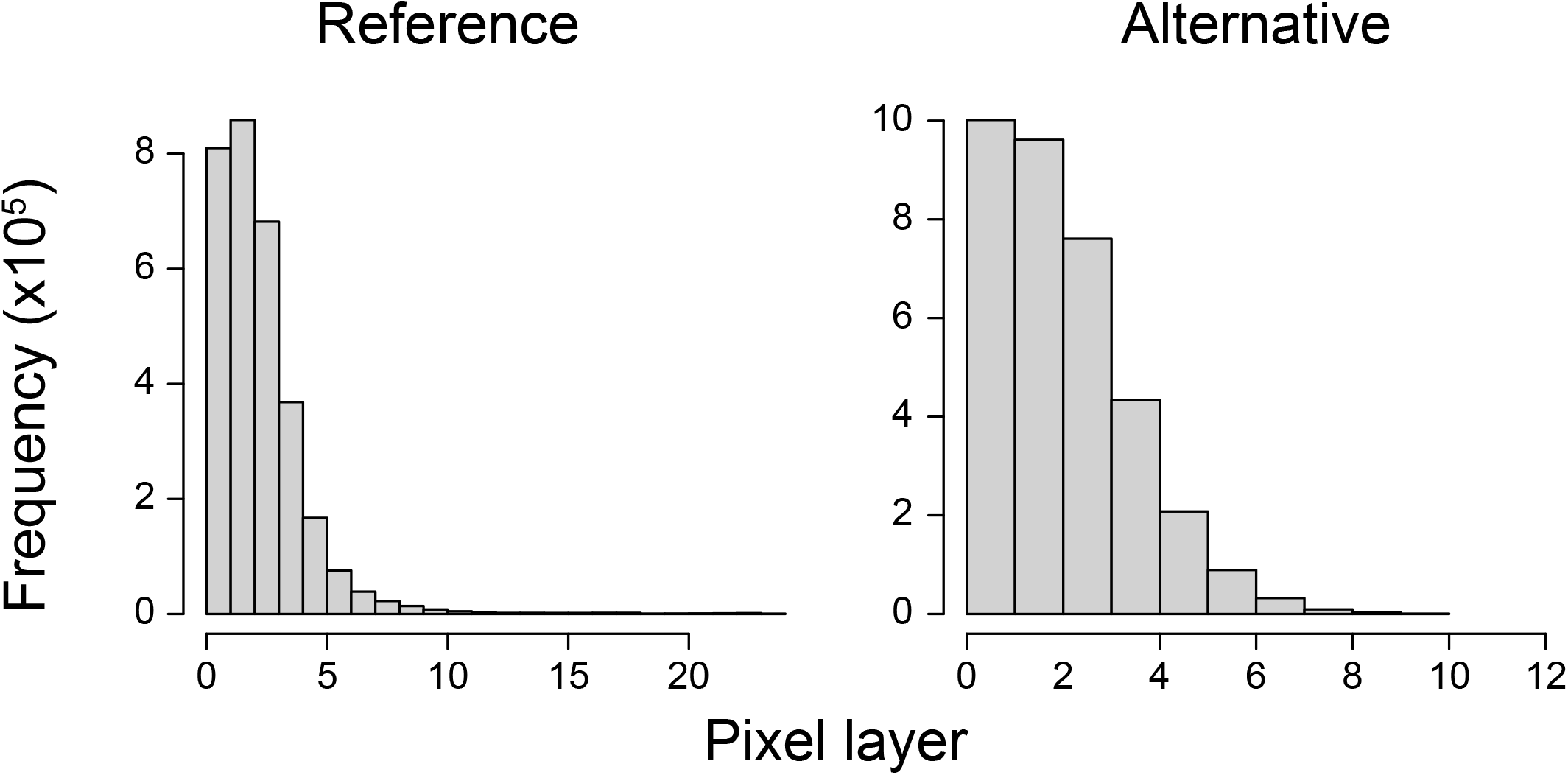
support to figure 7E. Distribution of pixel position within a cell.

